# Small-molecule activators of a bacterial signaling pathway inhibit virulence

**DOI:** 10.1101/2023.12.02.569726

**Authors:** Kathryn E. Mansour, Yunchuan Qi, Mingdi Yan, Olof Ramström, Gregory P. Priebe, Matthew M. Schaefers

## Abstract

The *Burkholderia* genus encompasses multiple human pathogens, including potential bioterrorism agents, that are often extensively antibiotic resistant. The FixLJ pathway in *Burkholderia* is a two-component system that regulates virulence. Previous work showed that *fixLJ* mutations arising during chronic infection confer increased virulence while decreasing the activity of the FixLJ pathway. We hypothesized that small-molecule activators of the FixLJ pathway could serve as anti-virulence therapies. Here, we developed a high-throughput assay that screened over 28,000 compounds and identified 11 that could specifically active the FixLJ pathway. Eight of these compounds, denoted *<u>B</u>urkholderia* <u>F</u>ix <u>A</u>ctivator (BFA) 1-8, inhibited the intracellular survival of *Burkholderia* in THP-1-dervived macrophages in a *fixLJ-*dependent manner without significant toxicity. One of the compounds, BFA1, inhibited the intracellular survival in macrophages of multiple *Burkholderia* species. Predictive modeling of the interaction of BFA1 with *Burkholderia* FixL suggests that BFA1 binds to the putative ATP/ADP binding pocket in the kinase domain, indicating a potential mechanism for pathway activation. These results indicate that small-molecule FixLJ pathway activators are promising anti-virulence agents for *Burkholderia* and define a new paradigm for antibacterial therapeutic discovery.

## Introduction

Members of the *Burkholderia* genus can cause serious, difficult to treat infections. Among *Burkholderia* species that cause serious infections in humans are the *Burkholderia cepacia* complex (BCC) and *Burkholderia pseudomallei*. The BCC is composed of approximately 20 species, many of which are significant pathogens for people with cystic fibrosis (CF) and chronic granulomatous disease (CGD)(1-3) and to an ever-growing group of hospitalized patients exposed to contaminated medications or medical devices.(4-17) *B. pseudomallei* causes melioidosis, a serious systemic infection that can include sepsis, pneumonia, fever, and abscesses. Melioidosis is life-threatening, and mortality can be as high is 50%.(18) *B. pseudomallei* is a potential bioterrorism agent and is classified as a Tier 1 agent by the U.S. Centers for Disease Control and Prevention. Although most commonly found in tropical and sub-tropical soil in South-East Asia and Australia, climate change is thought to have led to the recent isolation of *B. pseudomallei* from soil in Mississippi and Texas, and infections were linked to exposure to this soil.(19-21) *B. pseudomallei* also recently caused a cluster of 4 cases of melioidosis in the U.S. associated with contaminated aromatherapy spray.(22)

*Burkholderia* are intrinsically resistant to multiple antibiotic classes.(23) This resistance is mediated through alteration of antibiotic targets, decreased outer membrane permeability by modifications of LPS, decreased expression of porins, increased expression of antibiotic-inactivating enzymes, or increased production of efflux pumps.(23, 24) In a study of over 2,000 BCC isolates, more than 50% of the isolates were resistant to chloramphenicol, co-trimoxazole, ciprofloxacin, tetracycline, rifampin, and amoxicillin-clavulanate.(25) One study of 56 CF isolates of *B. dolosa* (a BCC member) found them to be nearly pan-resistant, with minocycline being the only active antibiotic (in just 29% of isolates).(26) Pan-resistance in outbreak strains of *B. cenocepacia* has also been reported.(27) While *B. pseudomallei* isolates are typically susceptible to β-lactam antibiotics, these antibiotics are often not effective at clearing the infection, and treatment can last for months.(18) Thus, novel compounds are needed for *Burkholderia* infections since the number of new prospects from traditional drug development is limited.

Our previous work focused on understanding the role of the *Burkholderia* FixLJ two-component system in pathogenicity. Two-component systems are one mechanism that bacteria use to sense and respond to their environment by modulating gene expression.(28, 29) We initially identified the *Burkholderia fixLJ* two-component system in bacterial whole-genome sequencing studies as being under strong positive selection during chronic *B. dolosa* or *B. multivorans* infection in people with CF.(30-32) The *fixLJ* system regulates ∼11% of the genome of *B. dolosa*(33) and is required for virulence. Our recent work showed that otherwise isogenic BCC constructs carrying evolved (late) *fixL* sequence variants are more virulent than constructs carrying ancestral (early) *fixL* sequence variants.(34) Interestingly, bacteria carrying these evolved *fixL* sequence variants have lower levels of FixLJ pathway activity, demonstrating that high levels of FixLJ pathway activity are detrimental to virulence.(34) These findings led us to the hypothesis that small-molecule activators of the *Burkholderia* FixLJ pathway could make the bacteria less virulent. In the current study, we describe a high-throughput screen that identified 11 novel activators of the *Burkholderia* FixLJ pathway. Eight of these compounds inhibited the virulence of *Burkholderia* in vitro in a *fixLJ*-dependent manner in intracellular survival assays using a macrophage cell line. The most active compound inhibited the virulence of multiple *Burkholderia* species, including *B. thailandensis*, a model organism for *B. pseudomallei*.

## Results

### High-Throughput Screen Identifies 84 Compounds that Activate the Burkholderia FixLJ Pathway

We developed and conducted a high-throughput screen to look for activators of the *Burkholderia* FixLJ pathway with the goal of identifying novel anti-virulence compounds. We modified an existing *fix* pathway reporter (33, 34) to express green fluorescent protein (GFP) instead of LacZ when the *fixK* promoter is activated by FixLJ. This construct was cloned into a mini-Tn7 based vector allowing for stable integration of the reporter into the *Burkholderia* chromosome without antibiotic selection.(35, 36) This reporter was then conjugated into the clinical CF isolate *B. multivorans* strain VC7102. For the screen, this reporter strain was incubated in 384-well plates in the presence of compounds or DMSO vehicle control, and both OD600 and GFP fluorescence were measured after overnight growth at 37°C (**Figure 1A**). In initial screens we evaluated a library of 640 FDA-approved compounds for their ability to induce the FixLJ pathway. Among this library were several antibiotics and other compounds that inhibited bacterial growth and GFP activity (lower left region of **Figure S1A**). We identified one compound, an anti-Parkinson’s disease drug called benserazide, that was able to induce GFP levels above vehicle (DMSO) treated wells (**Figure S1A**). We sought to confirm benserazide as a hit across two BCC species, and determine if it was *fixLJ-*specific, by using the same GFP reporter conjugated in *B. dolosa* strain AU0158 and its *fixLJ* deletion mutant. If benserazide specifically targeted FixLJ, there would not be an increase in fluorescence seen in the *fixLJ* deletion mutant when treated with benserazide. Benserazide was able to induce GFP response in a dose-dependent manner in both *B. multivorans* and *B. dolosa* (**Figure S1B**), but was it also induced a GFP response in the *fixLJ* deletion mutant, demonstrating that benserazide activates the GFP reporter in a *fixLJ* independent mechanism. Benserazide was subsequently used a positive fluorescence control in screening assays.

**Figure 1.**
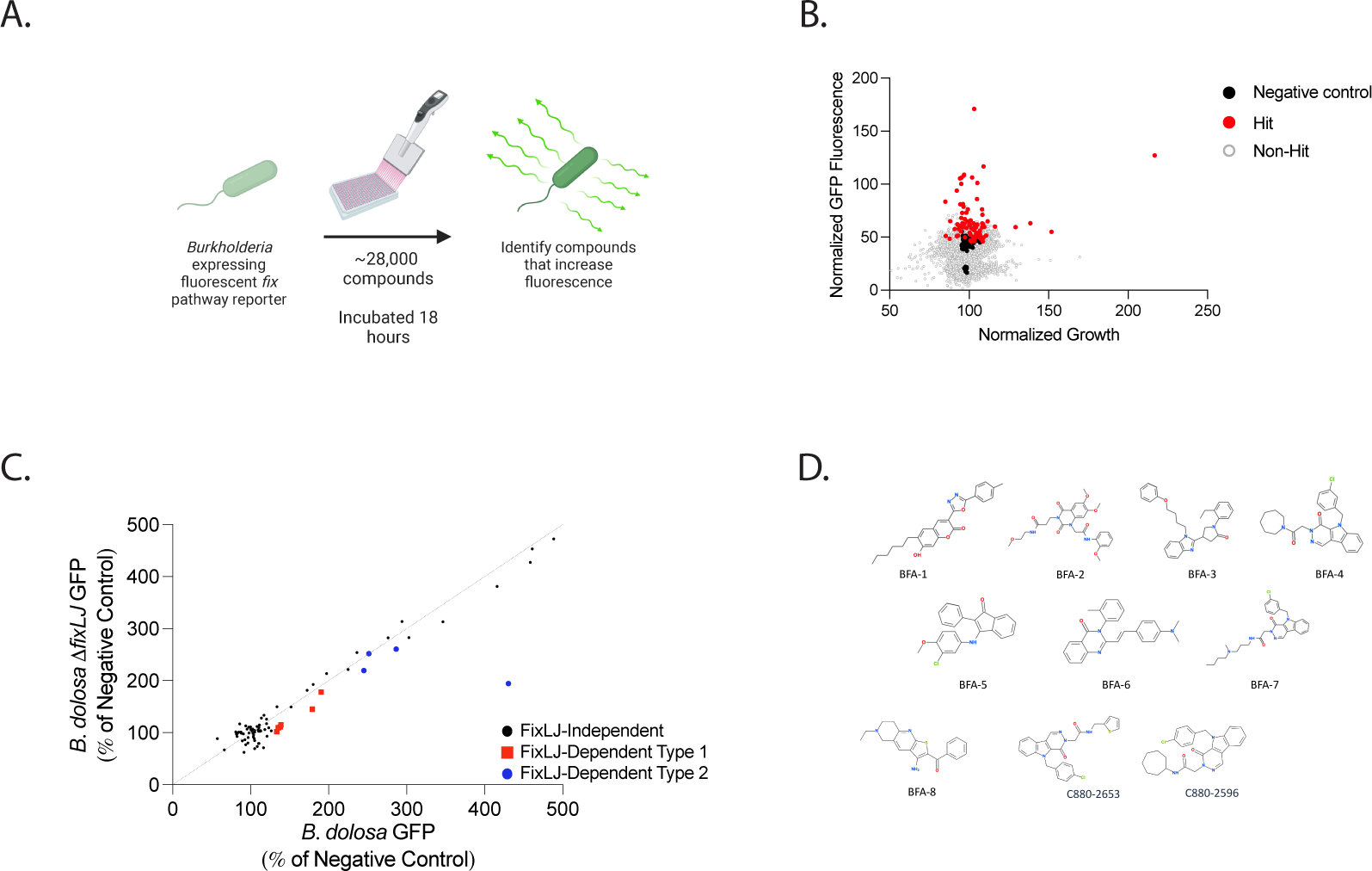
A high-throughput screen identifies 84 small molecules that activate the Burkholderia FixLJ pathway, 11 of which are *fixLJ-*specific. (A.) Schematic of the screen. Created with BioRender. (B.) Scatter plot from high-throughput screen of 28,100 compounds for activators of GFP activity in *B. multivorans* strain VC7102 carrying a GFP reporter for FixLJ pathway activity. GFP fluorescence and growth OD600 values are normalized based on plate-specific benserazide treated wells. Dots are averages of two replicate wells treated with same compound. Hits (red dots) have that GFP activity more than 3 standard deviations above the plate-specific mean negative control wells (DMSO, black dots). (C.) Scatter plot of 84 hits from primary screen chosen for follow-up assays measuring the GFP signal in *B. dolosa* and its *fixLJ* deletion mutant to assess dependence on FixLJ pathway. The GFP as percent of negative control (DMSO-treated) for each compound is plotted as GFP seen the *fixLJ* deletion mutant vs. parental strain. Type 1 hits (red) have that GFP activity more than 3 standard deviations above the plate-specific mean negative control wells in the parental strain, but not in the *fixLJ* deletion mutant. Type 2 hits (blue) are categorically greater hits in strength in the parental strain compared to the hit strength in the *fixLJ* deletion mutant. (D.) Structures of the 10 of the 11 *fixLJ-*dependent hits. These ten compounds were available from ChemDiv.

Next, 28,100 compounds were screened, with each assay plate including at least 1 column (16 wells) that contained DMSO (vehicle) alone as a negative control and at least 1 column (16 wells) that contained benserazide which served as a positive fluorescence control (**Figure 1**). Using benserazide as a positive fluorescence control and DMSO as a negative control, we were able to achieve Z’ factors ∼0.5, indicating the screening assay was technically robust.(37) We identified an additional 83 hits having mean fluorescence of the 2 replicate plates at least 3 standard deviations above the plate-specific negative control. Compounds were identified as weak hits if their increase in fluorescence was between 3-6 standard deviations above the negative control mean. Moderate hits had an increase in fluorescence between 6-9 standard deviations, and strong hits were at least 9 standard deviations above the mean negative control value (**Table 1**). These 83 compounds were “cherry-picked” from compound library plates, and their ability to induce GFP activity in *B. dolosa* strain AU0158 and its *fixLJ* deletion mutant was assessed to confirm FixLJ-specific activation of the GFP reporter (**Table S1**). We found that most compounds activated the GFP reporter to similar levels in the parental *B. dolosa* strain as in the *fixLJ* deletion mutant, indicating that these compounds were activating the reporter in a FixLJ-independent manner. We did find 7 compounds that activated the GFP reporter only in the parental *B. dolosa* strain, and we therefore classified these as type 1 hits (Red dots, **Figure 1C**). We also identified 4 compounds that were classified as hits in both the parental *B. dolosa* strain and the *fixLJ* deletion mutant, but were a lesser strength hit in the *fixLJ* deletion mutant compared the parental *B. dolosa*, so these were classified as type 2 hits (Blue dots, **Figure 1C**). It is important to note that these hits did not inhibit bacterial growth in the screen. In total we identified 11 compounds that were able to activate the FixLJ pathway (either type 1 or 2 hits), 10 of which were available commercially. The structures of these 10 compounds are depicted in **Figure 1**, and full chemical names are listed in Table S2.

**Table 1.** Number of hits from primary screen and number of hits confirmed to be *fixLJ-*specific.

| Type of Hit | # of Hits in Primary Screen | # of Confirmed *fixLJ*- Specific Hits |
| --- | --- | --- |
| Strong | 15 | 3 |
| Medium | 11 | 4 |
| Weak | 57 | 4 |

### Eight of the Small-molecule FixLJ Activators Inhibit Burkholderia Virulence in vitro

We tested the ability of the 10 hits to inhibit *B. dolosa* invasion of and/or intracellular survival within macrophages, which is a critical aspect of *Burkholderia* virulence.(38-40) Here, we used *B. dolosa* strain AU0158, a CF clinical isolate that also employed in the screen. In these assays, THP-1 cells are differentiated into macrophage-like cells using phorbol 12-myristate-13-acetate (PMA) and infected with *Burkholderia* (5-10 bacteria per macrophage) for two hours while being exposed to compound or vehicle (DMSO). Cells were washed and then treated with kanamycin (to kill extracellular bacteria) along with compound or DMSO for an additional 2-4 hours. Cells were washed again, lysed, and then colony forming units (CFU) were determined by serial dilution and plating (**Figure 2A**). Eight of the 10 compounds inhibited invasion/survival of *B. dolosa* in macrophages and were named *<u>B</u>urkholderia* <u>F</u>ix <u>A</u>ctivator (BFA) 1-8 (**Figure 2**). BFA1 inhibited *B. dolosa* virulence at concentrations as low as the lowest tested dose of 1.5 µM (**Figure 2B**) while the other 7 compounds inhibited virulence at concentrations between 6.25 and 12.5 µM (**Figure 2C-I**). BFA compounds 2-8 inhibited the invasion/survival of *B. dolosa* up to 50% at the highest dose of compound tested. BFA1 was able to inhibit invasion/survival of *B. dolosa* in macrophages by ∼75%. This reduction in the number of intracellular bacteria was *fixLJ-*specific, as the *B. dolosa fixLJ* deletion mutant, which already is less virulent than its parental strain as previously reported,(33) did not have significant reductions in intracellular bacteria when treated with any of the BFA compounds (**Figure 2**, red bars). Two of the 10 hits were not able to inhibit the invasion/survival of *B. dolosa* in macrophages (**Figure 2J and 2K**) and were not further evaluated.

**Figure 2.**
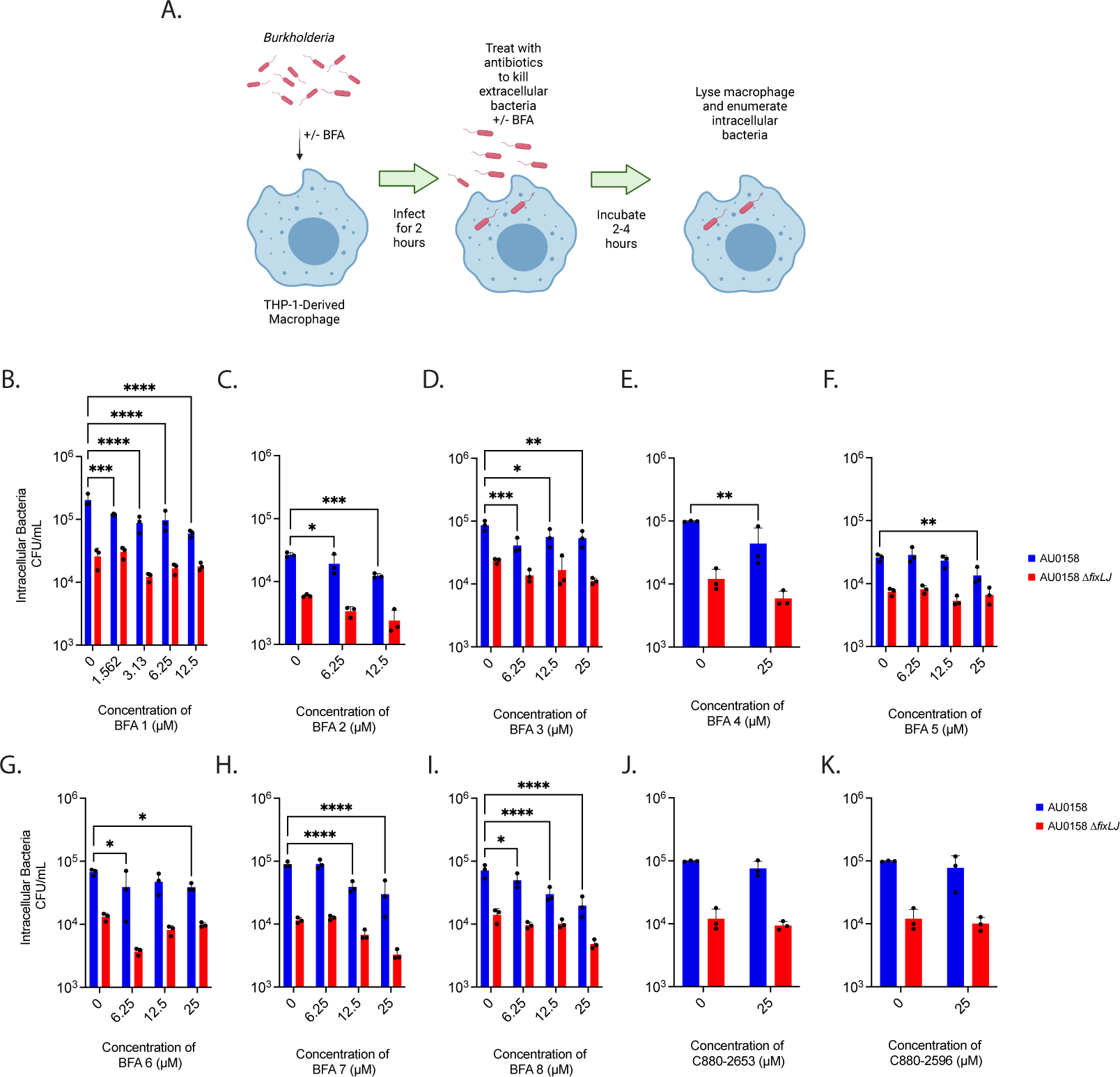
BFA (Burkholderia Fix Activator) compounds inhibit *B. dolosa* virulence in THP-1-derived macrophages in a *fixLJ*-specific manner. The intracellular survival (and/or uptake) of *B. dolosa* strain AU0158 in THP-1-derived human macrophages was measured using an antibiotic exclusion assay in the presence of varying concentrations BFA compounds. Created with BioRender. (A). The number of intracellular bacteria was determined by lysing the macrophages and enumerating CFU after incubation for 2 hours (B,D) or 4 hours (C,E,F,F,G,H,I,J,K). *, **, and *** denote *p*<0.05, 0.01, and 0.001, respectively, by two-way ANOVA with Dunnett’s multiple comparisons test using 0 µM (DMSO vehicle) as control.

We also measured the cytotoxicity of BFA compounds by measuring lactate dehydrogenase (LDH) release from a human lung epithelial cell line (A549) and from THP-1 derived macrophages using commercially available kits. As shown in **Figure S2**, we found that none of the compounds caused significant cytotoxicity after overnight exposure at any of the tested concentrations. These findings demonstrate that BFA compounds can inhibit the virulence of *Burkholderia* in a *fixLJ-*specific mechanism without significant toxicity *in vitro*.

In additional to measuring the ability of BFA compounds to inhibit the virulence of *B. dolosa*, we measured the ability of select BFA compounds to inhibit the invasion of and/or intracellular survival of other pathogenic *Burkholderia* species using the same THP-1-dervied macrophage infection model. We measured the ability of 7 of the 8 BFA compounds (25 µM) to inhibit invasion/intracellular survival of *B. cenocepacia, B. multivorans,* and *B. thailandensis*. *B. thailandensis* is a model for *B. pseudomallei* that does not require BSL3 facilities.(41) We were unable to obtain sufficient amounts of BFA4 for further analysis, so it was excluded. BFA1 inhibited the invasion/survival of all three additional *Burkholderia* species (**Figure 3A-3C**). All seven of the tested BFA compounds inhibited the invasion/survival of *B. multivorans* (**Figure 3A**). BFA1 was also able to inhibit the invasion/survival of *B. cenocepacia* (**Figure 3B**) and *B. thailandensis* (**Figure 3C**). BFA6 was also able to inhibit the invasion/survival of *B. thailandensis* (**Figure 3C**), while there was a trend towards significance for other BFA compounds to inhibit *B. thailandensis*. Since BFA1 had the highest activity, multiple doses were evaluated for their ability to inhibit invasion/survival of other *Burkholderia.* Doses of 6.25 µM of BFA1 inhibited the invasion/survival of *B. multivorans* (**Figure 3D**) and *B. thailandensis* (**Figure 3F**), while higher doses were needed to inhibit *B. cenocepacia* (**Figure 3E**).

**Figure 3.**
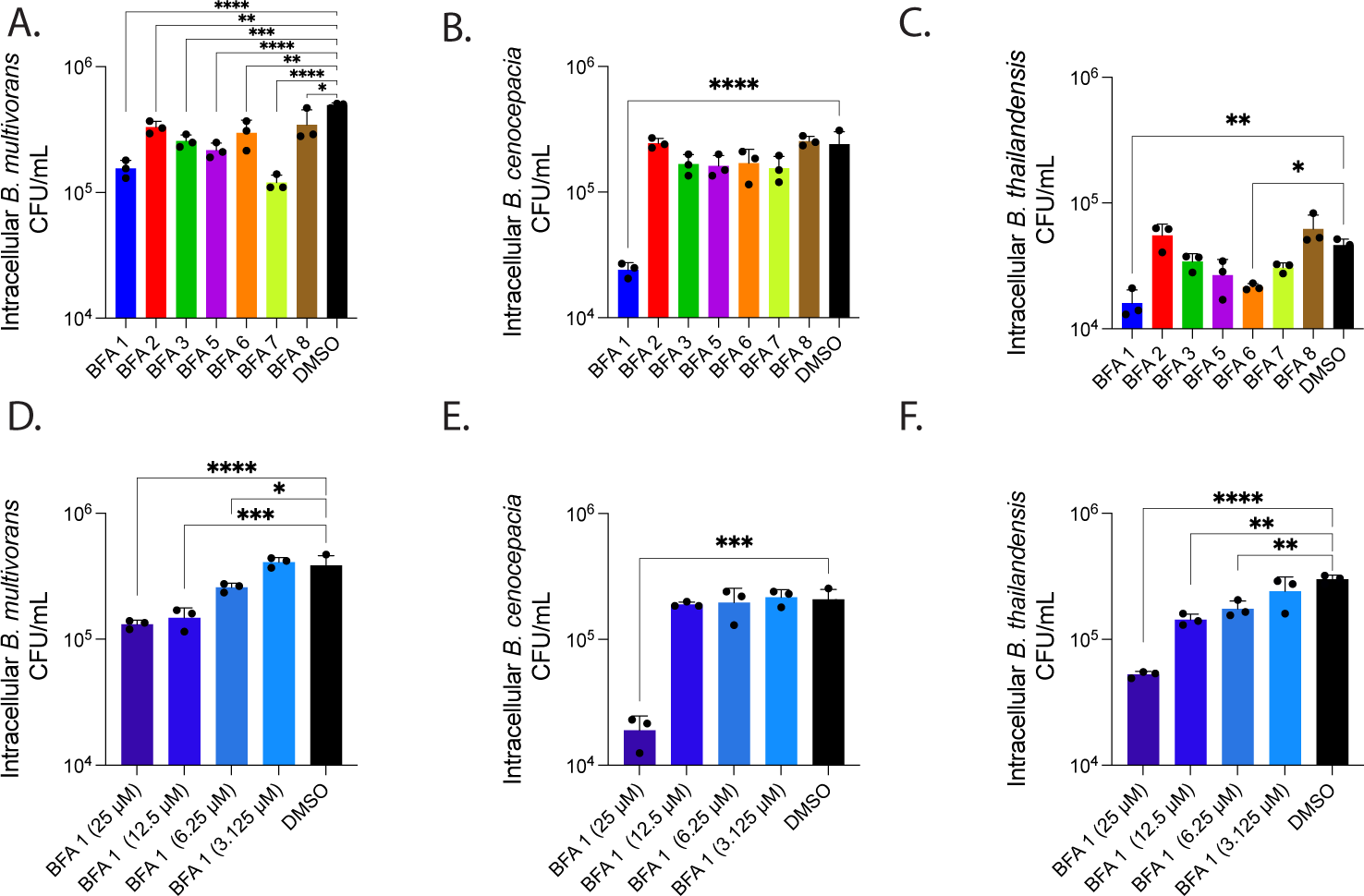
BFA compounds inhibit the virulence of multiple pathogenic *Burkholderia* species. THP-1-derived macrophages were infected with *B. multivorans* strain VC7102 (A&D), *B. cenocepacia* strain K56-2 (B&E), or *B. thailandensis* strain e264 (C&F) in the presence of 25 µM of BFA compounds (A-C) or a dose range of BFA1 (D-F). Intracellular bacteria were determined using antibiotic exclusion after 2 (F) or 4 (A-E) hour exposure to antibiotic. *P* value determined by ANOVA with Dunnett’s multiple comparisons test, *, **,***,**** denotes *p* value < 0.05,0.01, 0.001, 0.0001, respectively.

### In silico Docking Studies Demonstrate BFA1 Interaction with ATP/ADP-Binding Pocket of FixL

We performed docking studies with *B. dolosa* FixL (AlphaFold ID: A0A0D5J096) using AutoDockFR in the flexible residue mode.^(42)^ Initial AutoSite calculations predicted two major potential ligand-binding sites on the protein (**Figure S3A**, **Table 2**), of which the site with the highest affinity strongly resembled the ATP/ADP binding site of known histidine kinases, such as the structures from *Caulobacter vibrioides* (PDB ID: 5IDJ),(43) *Lactiplantibacillus plantarum* (4ZKI),(44) or *Thermotoga maritima* (6RH8).^(45)^ This site was thus chosen as the primary docking target, to which docking with ADP yielded a binding interaction similar to the mentioned histidine kinases, thereby validating the docking process (**Figure S3B**).

**Table 2.** Potential ligand-binding pocket parameters.

| Pocket | AutoSite score | Number of points | Radius of gyration | Buriedness |
| --- | --- | --- | --- | --- |
| 1 | 43.69 | 347 | 5.71 | 0.85 |
| 2 | 39.25 | 301 | 5.18 | 0.82 |

As expected, the predicted score for ADP binding to the second pocket was considerably lower. On the other hand, the binding scores for compound BFA1 were of similar magnitude for the two sites, in both cases better than for ADP (**Table S3**). For the ATP/ADP binding site, the docking analysis showed interactions between compound BFA1 and several residues in the binding pocket, with the coumarin oxygen atoms interacting with Gly777, Leu778, Thr769, and the oxadiazole oxygen interacting with Met776 (**Figure 4A, Figure S3C**). For the secondary binding site, the analysis revealed polar interactions between the ligand and residues Arg585, Asn779, and Ser783, as well as more lipophilic interactions with nonpolar residues (**Figure S3D**). Docking simulations were also carried out using HADDOCK 2.4,(46, 47) revealing similar binding trends (**Tables S3 and S4**). A potential effect may also arise from binding to the second pocket, although the results in this case indicate a lower degree of more specific polar interactions compared to the ADP/ATP binding site.

**Figure 4.**
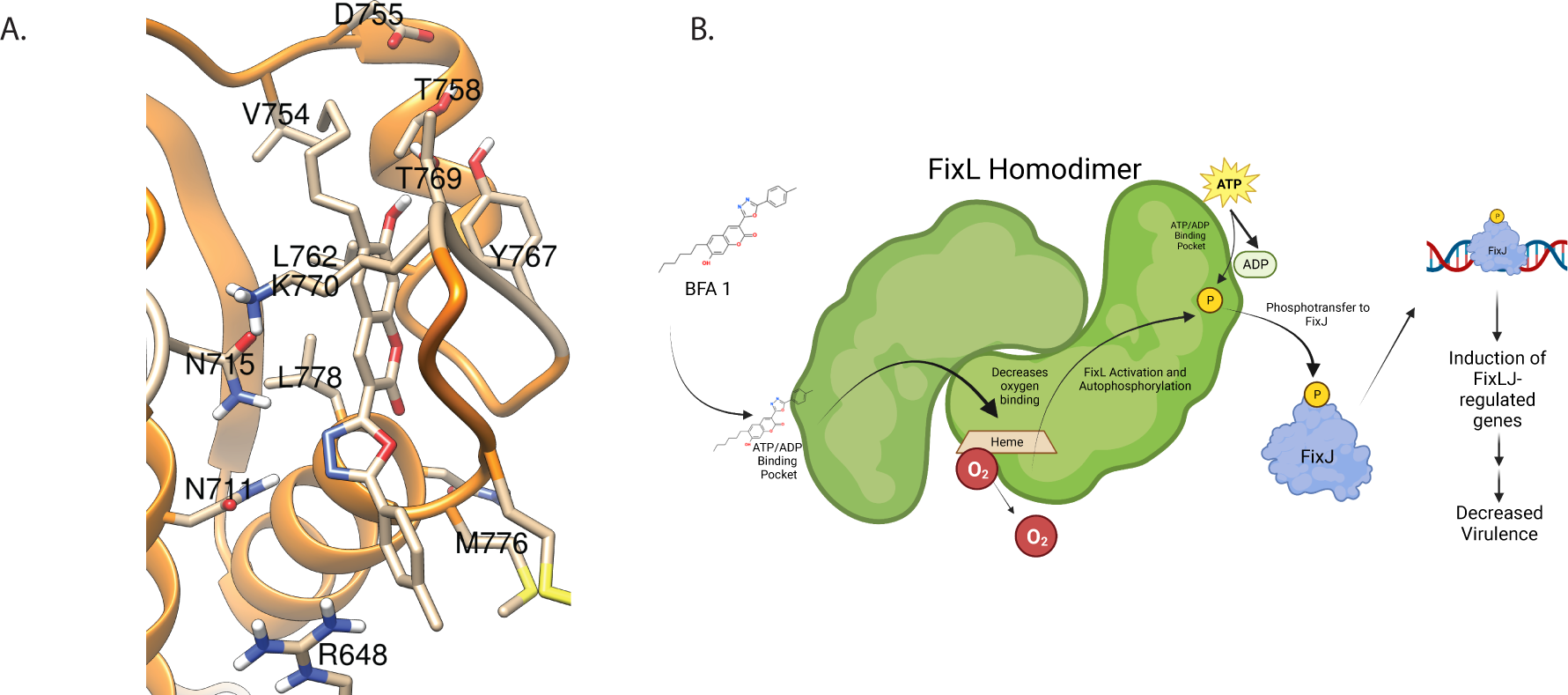
BFA1 is predicted to bind to FixL at the ATP/ADP-binding pocket of the histidine kinase domain. (A) Predicted binding of BFA1 to first binding pocket of *B. dolosa* strain AU0158 FixL using AutoDockFR. (B) Proposed mechanism of action for BFA1 activating *Burkholderia* FixLJ pathway. Created with BioRender.

## Discussion

In this study we developed a high-throughput screen to identify small molecules that activate the *Burkholderia* FixLJ pathway, with the goal of inhibiting the virulence of these pathogens. This approach was based on our previous work that found mutations within the *fixL* gene leading to lower FixLJ pathway activity were selected for during chronic infection in people with CF(30, 31) and correlated with periods of decline in lung function in people with CF.(34) These findings lead us to the hypothesis that a small molecule activating the FixLJ pathway could make *Burkholderia* less pathogenic, in a way coaxing the bacteria back to its soil existence where FixLJ activity is high.

Our high-throughput screen identified 8 compounds we call <u>B</u>urkholderia <u>F</u>ix <u>A</u>ctivators (BFA). These compounds previously had no known biological activity. All 8 of the BFA compounds inhibited the invasion and/or intracellular survival of *B. dolosa* in a *fixLJ-*specific manner (**Figure 2**) but had no impact on bacterial growth in rich media. BFA1 inhibited virulence at a lower concentration and at a greater magnitude than the other BFA compounds. The BFA compounds caused no significant cytotoxicity, measured by LDH release, after 24-hour incubation suggesting macrophage death plays a negligible role in the BFA activity in antibiotic exclusion assays. *Burkholderia* are known for their extensive antibiotic resistance that is often mediated by decreases in outer membrane permeability and increases in efflux pump activity.(23, 24) Since clinical isolates of *Burkholderia* were used in our assays that identified the BFA compounds, it is clear that the BFA compounds are able to overcome these intrinsic drug penetration obstacles that have hindered antibiotic development.

The *in silico* predicted binding of BFA 1 to the ATP/ADP binding pocket of FixL with a higher affinity than ADP is, on first review, counterintuitive for a molecule that increases FixL activity. However, for rhizobial FixL, ADP binding has been shown to decrease the affinity of FixL for oxygen, which results in activation of FixL.(48) This activation is related to the homodimer properties of FixL(49) whereby ADP that is formed as part of the autophosphorylation of FixL binds to FixL, which decreases the binding affinity to oxygen in the other FixL molecule of the homodimer. This, in turn, allows for a positive feedback resulting in increased FixL activation.(48) Thus, BFA1 binding to FixL at this same pocket is predicted to increase FixL activation as a result of decreased binding affinity for oxygen. These predictions are supported by findings that single amino acid changes of the asparagine residue (403) in the ADP-binding site of rhizobial FixL resulted in no change in oxygen affinity in response to ADP binding, demonstrating the importance of this residue in ADP binding.(48) The homologous asparagine in the predicted ADP-binding site of *Burkholderia* FixL is at amino acid 715 and is predicted to interact with or be adjacent to BFA1 or ADP binding (**Figure 4** and **S3**). The predicted mechanism of action is depicted in **Figure 4F**, where BFA1 binds to FixL via the ATP/ADP binding pocket on the protein, which causes a decrease in binding affinity to oxygen and, in turn, activates FixL. FixL autophosphorylates, then transfers the phosphate group to the response regulator FixJ. Phosphorylated FixJ then binds to DNA and turns on transcription of target genes that are part of the FixLJ regulon resulting in a gene expression profile making the bacteria less virulent.

Our method of using small-molecule activators of a pathway to inhibit the virulence of antibiotic-resistant pathogens is a novel approach for the development of new antibacterial therapies. Most of the other limited number of studies investigating two-component systems as drug targets have focused on two-component systems that are involved in quorum sensing pathways and other pathways required for bacterial growth.(28, 50-53) Other groups have identified compounds that <u>inhibit</u> bacterial two-component systems to make pathogens less virulent,(54-59) highlighting that two-component systems can be targeted in multiple ways to make a pathogen less virulent. By targeting bacterial virulence rather than bacterial growth, the emergence of resistance to therapies will be slower to occur.(60) We expect that resistance to our lead compound, BFA1, will be slow to develop since the amino acids of FixL predicted to be involved in binding to BFA1 are outside the FixL domains where mutations are seen during chronic infection.(30, 31) In conclusion, our results show that small-molecule activators of the *Burkholderia* FixLJ pathway are a promising new anti-virulence approach.

## Methods

### Bacterial strains, plasmids, cell lines, and growth conditions

Bacterial strains used and generated in this study are listed in **Table 3**. For the generation of the reporter strain, BCC and *E. coli* were grown on LB plates or in LB medium and supplemented with following additives: ampicillin (100 μg/mL), chloramphenicol (20 μg/mL), trimethoprim (100 μg/mL for *E. coli,* 1 mg/mL for BCC), gentamicin (50 μg/mL). For some experiments, trypticase soy broth (TSB) or trypticase soy agar (TSA) was used for growth of *Burkholderia*. Human monocyte line THP-1 and human lung epithelial cell line A549 were obtained from ATCC and grown at 37°C with 5% CO_2_. THP-1 cells were cultured in RPMI-1640 medium containing 2 mM L-glutamine, 10 mM HEPES, 1 mM sodium pyruvate, 4500 mg/L glucose, and 1500 mg/L sodium bicarbonate, supplemented with 10% heat-inactivated fetal calf serum (FCS, Gibco) and 0.05 mM 2-mercaptoethanol. A459 cells were grown in RPMI 1640 with L-glutamine and 10% heat-inactivated fetal calf serum (FCS, Gibco). Penicillin and streptomycin were added for routine culture but were removed the day before and during experiments.

**Table 3.** Strains used in this study.

|  | Notes | Source |
| --- | --- | --- |
| *E.coli* |  |  |
| NEB 5-alpha Competent *E. coli* | DH5α derivative cloning strain | NEB |
| *Burkholderia* |  |  |
| *B. dolosa* AU0158 | Clinical isolate | John<br>LiPuma |
| *B. dolosa* AU0158 Δ*fixLJ* | Clean, unmarked *fixLJ* deletion mutant | (33) |
| *B. multivorans* VC7102 | Clinical isolate | (32) |
| *B. multivorans* VC7102 Fix-GFP Reporter | *B. multivorans* VC7102 + Fix-GFP reporter | This study |
| *B. dolosa* AU0158 Fix-GFP Reporter | *B. dolosa* AU0158 + Fix-GFP reporter integrated at *attTn7* site downstream of AK34_4894 | This study |
| *B. dolosa* AU0158 $\Delta fixLJ$ Fix-GFP Reporter | *B. dolosa* $\Delta fixLJ$ + Fix-GFP reporter integrated at *attTn7* site downstream of AK34_4894 | This study |
| *B. thailandensis* strain e264 | Isolate from rice field in Thailand, model for *Burkholderia pseudomallei* | ATCC |
| *B. cenocepacia* K56-2 | Clinical Isolate | Joanna Goldberg |

### Genetic Manipulations and Strain Construction

All plasmids used and generated in this study are listed in **Table 4**. To generate the GFP reporter of FixLJ pathway activity (Fix-GFP reporter), we cloned the first 23 bp of *fixK* and the immediate 243 bp upstream of the start codon from pfixK-reporter(33) in-frame with eGFP gene from pIN301 into the multiple cloning site of pUC18-mini-Tn7-Tp. Use of a mini-Tn7 vector allows for stable chromosomally integration at an *att*Tn7 site.(36, 61) The plasmid was transformed into NEB 5-alpha competent *E. coli,* and the sequence of the plasmid was confirmed using PCR and Sanger sequencing. The Fix-GFP reporter was conjugated into *B. dolosa* strain AU0158, the AU0158 *fixLJ* deletion mutant, and *B. multivorans* strain VC7102 with pRK2013 and pTNS3 using published procedures.(33) Conjugants were selected for by plating on LB agar containing trimethoprim (1 mg/mL) and gentamicin (50 μg/mL). Insertions into the *att*Tn7 site downstream of AK34_4894 was confirmed by PCR of *B. dolosa* strains as previously published.(33, 34)

**Table 4.** Plasmids used in this study.

|  | Notes | Source |
| --- | --- | --- |
| pTNS3 | Amp<sup>R</sup>, helper plasmid for mini-Tn7 integration into *attTn7* site | (35) |
| pRK2013 | Km<sup>R</sup> conjugation helper | (61) |
| pUC18T-mini-Tn7T-Tp | Amp<sup>R</sup>, Tp<sup>R</sup> on mini-Tn7T; mobilizable | (61) |
| pfixK-reporter | pSCrhaB2 carrying *B. dolosa fixK* promoter -*lacZ* fusion, Tp<sup>R</sup> | (33) |
| pIN301 | eGFP, CmR | Annette Vergunst |
| Fix-GFP reporter | *fixK* promoter -GFP fusion in pUC18T-mini-Tn7T-Tp | This study |

### Small Molecule High Throughput Screens

Small-molecule screens were conducted at Institute of Chemistry and Cell Biology (ICCB) - Longwood Screening Facility at Harvard Medical School. *B. multivorans* strain VC7102, with Fix-GFP reporter, was grown overnight in TSB at 37°C with shaking and diluted 1:100 in fresh TSB the day of the assay. 30 µL per well was added into a black clear-bottom 384-well plate using Thermo Multidrop Combi pipette. 30 µL per well of TSB was also added into each well containing a test compound using the Combi pipette. Wells that served as negative controls contained 30 µL of TSB plus DMSO (0.078% v/v in TSB, final concentration 0.039%). For positive fluorescence control wells, we added 30 µL of TSB containing 78 µM of benserazide (final concentration 39 µM). 300 nL solutions of each compound in DMSO were pin-transferred to each plate using an Epson Compound Transfer Robot from a compound library plate. For most compound library plates the stock concentration was 10 nM, but some plates had different concentrations typically 1-10 mM, details are posted in screen data deposited in PubChem. For every assay, 2 replicate assay plates were set up. Initial OD600 and GFP fluorescence were measured using PerkinElmer EnVision with Photometric 600 filter (600 nm, 8 nm bandpass), FITC 535 (Excitation filter FITC 585, Emission Filter FITC 535). Plates were stacked 5 high, covered with lids, and incubated at 37 °C overnight (∼18 h). The following day, assay plates were read using the PerkinElmer EnVision as above. For each well, the initial GFP fluorescence intensity values were subtracted from overnight GFP fluorescence intensity values to calculate the ΔGFP for each well. The average ΔGFP was calculated by averaging the 2 replicate wells for each library well. The mean and standard deviation for ΔGFP of the negative control wells on the replicate plates was calculated. A compound was determined to be a strong hit if the average ΔGFP was more than 9 standard deviations above mean ΔGFP for the negative control wells on the 2 replicate plates. A compound was determined to be a moderate hit if the average ΔGFP was more than 6 standard deviations above mean ΔGFP for the negative control, and a compound was determined to be a weak hit if the average ΔGFP was more than 3 standard deviations above mean ΔGFP for the negative control for its respective plate. Subsequent analysis of data from the primary screen identified 3 additional hits that were not identified in first analysis and were not further evaluated. For “cherry-pick” studies, overnight cultures of *B. multivorans* strain VC7102*, B. dolosa* strain AU0158, and *B. dolosa* strain AU0158 *fixLJ* deletion mutant were diluted and plated in wells of a 384-well plate. 300 nL of selected compounds were plated into wells using a HP D300e liquid dispenser so that 2 wells of each bacterial strain were treated with compound. Plates were incubated, and the OD600 and GFP fluorescence intensity were measured as described above.

### Bacterial invasion assays

The ability of BFA compounds to inhibit the uptake of and/or intracellular survival of *Burkholderia* into THP-1-dervived macrophages was determined using published protocols.(33, 34) Human THP-1 monocytes were differentiated into macrophages by seeding 1 mL into 24-well plates at 7.5x10^5^ cells/mL with 200 nM phorbol 12-myristate 13-acetate (PMA). Log-phase *Burkholderia* were grown and washed in RPMI containing 10% heat-inactivated FCS three times and diluted to ∼2x10^6^ CFU/mL and mixed with BFA compounds or DMSO (vehicle control). 1 mL/ well (MOI of ∼10:1) of the bacterial suspension was used to infect THP-1 derived macrophages. Plates were spun at 500 g for 5 minutes to synchronize infection and then incubated for 2 hours at 37°C with 5% CO_2_. To determine the number of intracellular bacteria, separate infected wells were washed two times with PBS and then incubated with RPMI plus 10% heat-inactivated FCS containing BFA or DMSO (vehicle control) with kanamycin (1 mg/mL) or kanamycin plus ceftazidime (1 mg/mL each for *B. cenocepacia*) for 2-4 hours. Monolayers were washed three times with PBS, lysed with 1% Triton-X100, serially diluted, and plated to enumerate the number of bacteria.

### LDH release assay

A549 cells were grown to confluence in 96-well plates. Human THP-1 monocytes were differentiated into macrophages by 72-hour PMA treatment and seeded into 96-well plates at density of 7.5x10^4^ cells/well. A549 cells and THP-1-deried macrophages were then treated with BFA (0-25 µM in DMSO) for 24 hours. LDH release was measured within supernatants using CytoTox 96® Non-Radioactive Cytotoxicity Assay (Promega) per manufacture’s protocol. For a positive control, untreated wells were incubated with 10X lysis buffer (provided with kit) during the last 30 minutes. Percent cytotoxicity was determined relative to maximum LDH release from cells treated with lysis buffer.

### In silico docking studies

Docking studies were performed using AutoDockFR Suite 1.0 in the flexible residue mode. ^(42)^ Calculations were performed with 8 independent searches, each of which with 50 genetic algorithm evolutions associated with 2×10⁶ evaluations of the scoring function. Potential binding sites were identified by AutoSite 1.0. Residues Asn715, Tyr767, Ser768, Thr769, and Lys770 were set as flexible for pocket 1 and residues Arg585, Asn779, Ser783 for pocket 2. Docking simulations were also carried out using HADDOCK 2.4,(46, 47) using the default settings for small molecule-protein docking with RMSD-based clustering. The regions of the two binding sites identified by ADFR were investigated using corresponding active residues (Asn715, Tyr767, Ser768, Thr769, Lys770 at pocket 1; Arg585, Asn779, Ser783 at pocket 2). 1000 structures were generated initially and 200 clusters were screened out after refinement. All docking results were visualized by UCSF Chimera 1.17.3 and LigPlot+ 2.2.(62, 63)

## Data Availability

Data from the high-throughput screen are deposited to PubChem AID 1918990.

## Acknowledgements

We would like to thank the staff of the Institute of Chemistry and Cell Biology (ICCB) - Longwood Screening Facility at Harvard Medical School for their assistance with the screening assays. This work was funded by National Institutes of Health (R21AI159211 to MMS), the Department of Anesthesiology, Critical Care and Pain Medicine at Boston Children’s Hospital (Trailblazer Award and Transition to Independence Award to MMS, no numbers), and the Cystic Fibrosis Foundation (PRIEBE13I0 to GPP). OR and MY thank the University of Massachusetts Lowell for financial support (No Number).

## Author Contributions

GPP and MMS conceived of the idea for the study. KEM, YQ, MY, OR and MMS conducted the experiments. MY, OR, GPP and MMS acquired funding. MY, OR, GPP, and MMS supervised. KEM and MMS drafted original draft, all authors revised and approved final version.

