## Supplementary material for "Small-molecule activators of a bacterial signaling pathway inhibit virulence": Figure S1

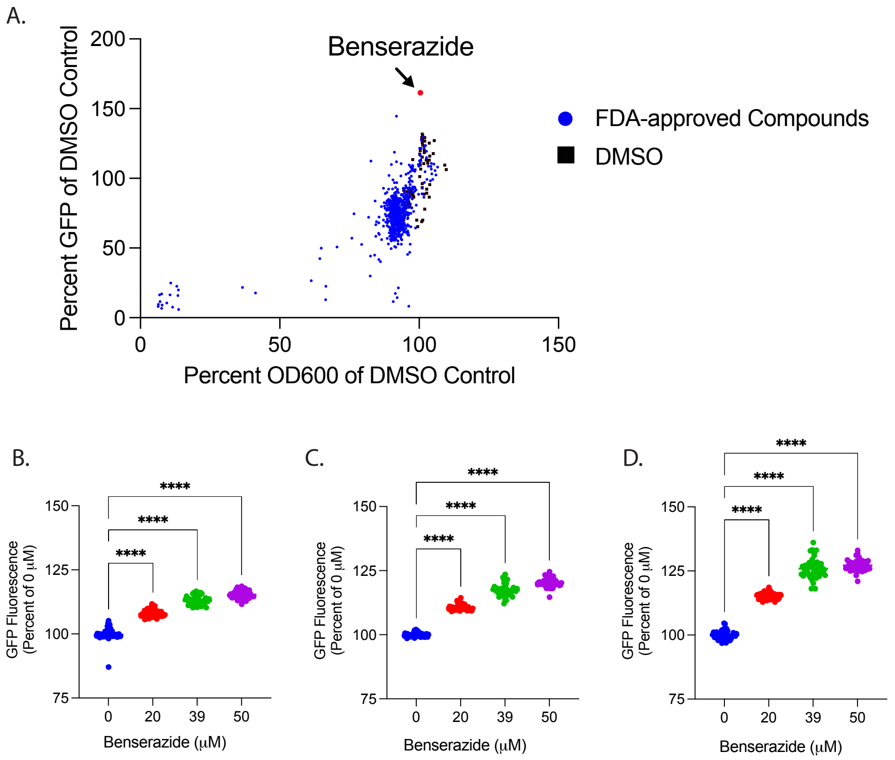


**Figure S1. Identification of benserazide as *fixLJ*-independent activator of the FixLJ pathway for use as positive fluorescence control.** (A.) Scatter plot of the impact of 640 FDA-approved compounds on GFP activity in *B. multivorans* strain VC7102 carrying GFP reporter for FixLJ pathway activity. Each blue dot depicts the mean of replicate plates containing the same compound. The GFP value was the ΔGFP value as described in Methods. Both GFP and OD600 are plotted as a % of plate specific-DMSO vehicle (negative) controls depicted in black dots. Benserazide (Red dot) was the only compound for which GFP activity was more than 3 standard deviations above the mean of the negative control (DMSO). Dose-dependent benserazide activation of the Fix-GFP reporter plasmid was measured in *B. multivorans* strain VC7102 (B), *B. dolosa* strain AU0158 (C), or *B. dolosa* strain AU0158 Δ*fixLJ* (D) was measured and plotted as a percent of 0 μM (DMSO only). **** denotes $p<0.0001$ by ANOVA with Dunnett's Multiple comparison test using 0 μM as control.
