## Supplementary material for "Small-molecule activators of a bacterial signaling pathway inhibit virulence": Figure S2

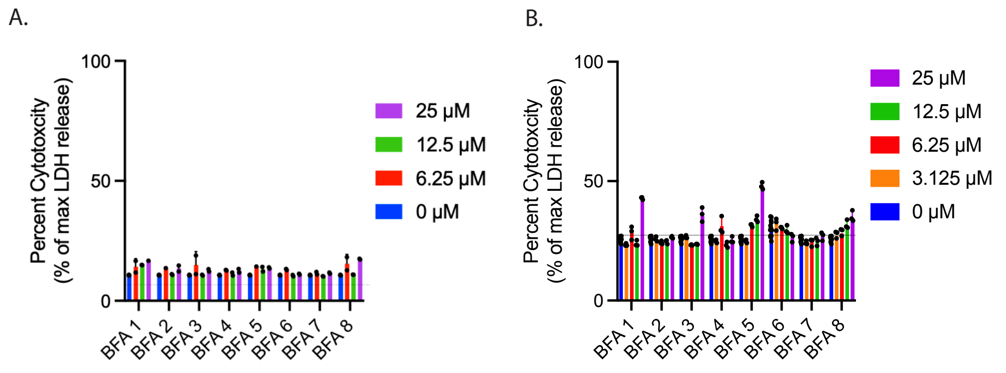


**Figure S2. BFA compounds are not cytotoxic to human cell lines.** Varying concentrations of compounds were incubated with A549 epithelial cells (A) or THP-1-dervived macrophages (B) for 16 hours, when LDH release was measured and plotted as maximum LDH release when cells were treated with detergent to completely lyse cells. Dotted lines denote untreated cells.
