## Supplementary material for "Small-molecule activators of a bacterial signaling pathway inhibit virulence": Figure S3

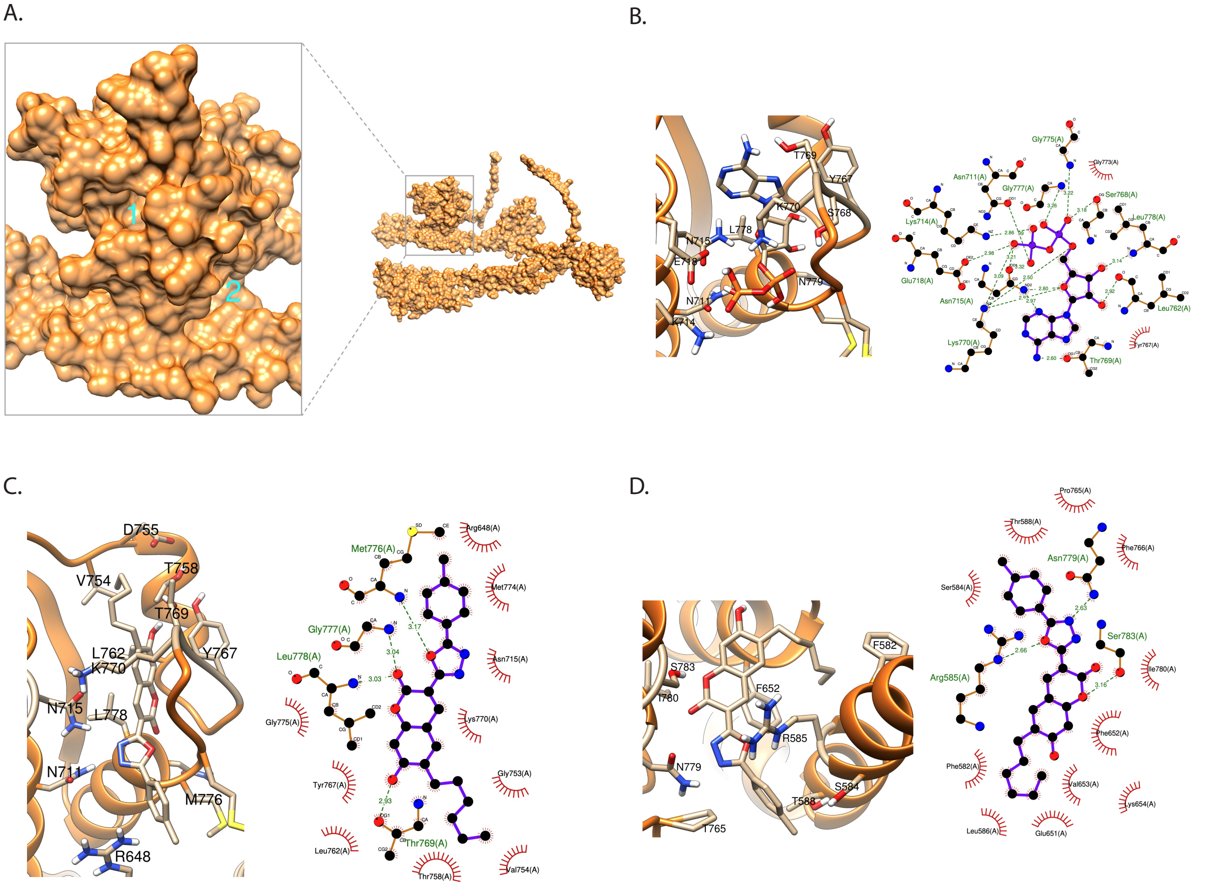


**Figure S3. In silica binding of BFA1 and ADP to *B. dolosa* FixL.** (A) Predicted FixL ligand binding pockets with highest scores. (B) ADP/FixL complex at ATP/ADP binding site predicted by ADFR (left) and interaction diagram (right). (C) BFA1/FixL complex at ATP/ADP binding site predicted by ADFR (left) and interaction diagram (right). (D) BFA1/FixL complex at pocket 2 predicted by ADFR (left) and interaction diagram (right).
