## Supplementary material for "Small-molecule activators of a bacterial signaling pathway inhibit virulence": Table S3

**Table S3.** Ligand/protein clustering information predicted by ADFR

| **Pocket** | **Ligand** | **Aﬃnity (kcal/mol)** | **Clust. RMSD** | **Ref. RMSD** | **Clust. size** | **RMSD SD** | **Energy SD** |
| --- | --- | --- | --- | --- | --- | --- | --- |
| 1 | ADP | -9.4 | 0 | -1 | 21 | 0.5 | 0.7 |
| 2 | ADP | -6.9 | 0 | -1 | 5 | 0.7 | 0.3 |
| 1 | BFA1 | -12.6 | 0 | -1 | 4 | 0.8 | 0.4 |
| 2 | BFA1 | -12.4 | 0 | -1 | 11 | 0.6 | 0.5 |
