## Supplementary material for "Small-molecule activators of a bacterial signaling pathway inhibit virulence": Table S4

**Table S4**: Ligand/protein clustering information predicted by HADDOCK.

| **Pocket** | **Ligand** | **HADDOCK score** | **No. clusters** |
| --- | --- | --- | --- |
| 1 | ADP | -82.708 | 64 |
| 2 | ADP | -55.213 | 112 |
| 1 | BFA1 | -48.647 | 16 |
| 2 | BFA1 | -46.693 | 76 |
